# Unfolded protein response signaling promotes myeloid cell production and cooperates with oncogenic mutation

**DOI:** 10.1101/2025.09.07.674755

**Authors:** Hyunjoo Choi, Sang-Eun Jung, Hyojung Paik, Yesenia Earth, Jiye Yi, Maggie J. Cox, Stephen M. Sykes, Stephen T. Oh, Yoon-A Kang

## Abstract

Unfolded protein response (UPR) promotes protein homeostasis under endoplasmic reticulum stress. UPR signaling has numerous functions in metabolism, cancer, immunology, and neurodegenerative diseases. Recent studies also showed that UPR signaling has important roles in hematopoietic stem and progenitor cell biology. However, whether UPR signaling regulates hematopoietic lineage fate decision remains elusive. Here, we found that FcγR^−^ MPP3 generates erythroid lineage and *Jak2^V617F^* mutation leads to overproduction of erythroid cells by expanding FcγR^−^ MPP3. We showed that UPR signaling increases myeloid cell production through promoting FcγR^−^ MPP3 transition to granulocyte/macrophage progenitor producing FcγR^+^ MPP3 at the expense of erythroid lineage via the XBP1 pathway. Under a disease condition, UPR signaling cooperates with *Jak2^V617F^* mutation and exacerbates disease phenotype in a mouse model of polycythemia vera (PV) through the ATF4 pathway. Activation of UPR signaling also increased myeloid output in healthy donor bone marrow MPP cells while skewing the output towards erythroid lineage in PV patient bone marrow MPP cells. Together, our results identify a novel function of UPR signaling in hematopoietic lineage specification and provide critical insights into targeting UPR signaling in hematological malignancies.

**Key points:**

- UPR signaling promotes myeloid cell production at the expense of erythroid lineage in steady state.
- UPR signaling collaborates with *Jak2^V617F^* mutation and increases red blood cell production.

## Introduction

Myelopoiesis is a dynamic process starting from hematopoietic stem cells (HSCs) and multipotent progenitor (MPP) populations that incorporates various environment cues and produces myeloid cells to adapt organismal needs^1^. Activation of HSCs and remodeling of MPP compartment such as expansion of myeloid-biased MPP2 and MPP3 and myeloid-reprogramming of lymphoid-biased MPP4 lead to granulocyte/macrophage progenitors (GMP) expansion followed by mature myeloid cell production^2–4^. In particular, secretory MPP3 subset, ER^high^/FcγR^+^ MPP3 expansion, which is triggered by high Wnt and low Notch activity, directly amplifies myelopoiesis by secreting pro-myeloid cytokines^5,6^. Pro-inflammatory cytokines like IL-6, IL-1, or TNFα also act on multiple steps during this process to boost myeloid cell production^2,4,7,8^. Given the significant contribution of MPP to blood production and the functional consequences of lineage priming at the MPP levels^9,10^, what regulates the MPP compartment, especially the myeloid-biased MPP holds tremendous implications in controlling myelopoiesis.

The unfolded protein response (UPR), which was initially identified as a response to the abnormal accumulation of unfolded or misfolded proteins, is a signal transduction network comprised of three central pathways; the IRE1α-XBP1, PERK-ATF4, and ATF6 pathways^11^. Aberrant UPR signaling has been implicated in various pathophysiologies including inflammation, metabolic disease, neurodegenerative diseases, and cancer^11^. Recent studies also show that the UPR signaling possesses diverse functions beyond protein homeostasis^11^. In the hematopoietic system, UPR preserves HSC pool integrity under stress condition^12^. IRE1α–XBP1 axis maintains HSC self-renewal and protects hematopoietic stem and progenitor cells (HSPCs) from myeloid leukemogenesis^13,14^. XBP1 controls cytokine secretion in macrophages, B cell differentiation, dendritic cell development, and platelet aggregation^15–19^. ATF4 deficiency reduces the number of functional HSC in the fetal liver and impairs HSC function as well as causes erythroid differentiation defects in adult mice^20–23^. Although UPR regulates many aspects of hematopoiesis, whether UPR signaling controls lineage fate decision in HSPCs is poorly understood. Here, we report that UPR signaling promotes myeloid production at the cost of erythroid lineage and potentiates the pathophysiology of *Jak2^V617F^*mutation under a disease condition, identifying the new regulatory function of UPR signaling in hematopoiesis.

## Methods

### Mice

All animal experiments were conducted at Washington University in accordance with IACUC protocols. CD45.2 C57BL/6J (000664), CD45.1 C57BL/6-BoyJ (002014) and B6.Cg-Tg(Mx1-cre)1Cgn/J (003556) mice were purchased from the Jackson Laboratory. Eight- to twelve-week-old CD45.2 C57BL/6J and CD45.1 C57BL/6-BoyJ were used. Cre driven *Jak2^V617F^*knock-in mice^24^ were used 12 weeks post-induction for *UBC-* and *Mx1-Cre* and when they reached at least 12 weeks old for *Vav-Cre*. *Mx1-Cre* driven *Xbp1* floxed mice^25^ were used 8 weeks post-deletion. Respective littermates were used as controls.

### Patient and healthy donor samples

Patient bone marrow samples were obtained according to a protocol approved by the Washington University Human Studies Committee (WU no. 01-1014). All patients previously provided consent to have samples banked and were not newly recruited for this study. Healthy donor bone marrow samples were purchased from StemCell Technologies (70001) or Lonza (2M-125C). Patient sample list used in this study is provided in Supplementary Table 4.

### Flow cytometry

Staining of hematopoietic cells was performed as previously described^5^. Antibody list used in this study is available in Supplemental Table 5. MPP2 is isolated as Lin^−^/Sca-1^+^/c-Kit^+^/Flk2^−^/CD150^+^/CD48^+^; FcγR^−^ MPP3 as Lin^−^/Sca-1^+^/c-Kit^+^/Flk2^−^/CD150^−^/CD48^+^/FcγR^−^; FcγR^+^ MPP3 as Lin^−^/Sca-1^+^/c-Kit^+^/Flk2^−^/CD150^−^/CD48^+^/FcγR^+^; MPP4 as Lin^−^/Sca-1^+^/c-Kit^+^/Flk2^+^/CD150^−^/CD48^+^. For short-term *in vitro* culture of FcγR^−^ MPP3 with tunicamycin, cells were stained with Sca-1-BV421, c-Kit-APC-Cy7, CD48-A647, CD150-PE, and FcγR-PE-Cy7. For short-term *in vivo* differentiation assays, c-Kit-enriched recipient BM cells were stained with lineage cocktails (CD3, CD4, CD5, CD8, CD11b, B220, Gr1, Ter119), CD45.1-PE, CD45.2-FITC, Sca-1-BV421, c-Kit-APC-Cy7, CD48-A647, CD150-PE, FcγR-PE-Cy7 and CD34-Bio followed by SA-BV605. Stained cells were re-suspended in staining media containing 1 µg/ml propidium iodide (PI) for dead cell exclusion.

### In vitro assays

For short-term *in vitro* culture, cells were grown in 200 µl base media consisting of Iscove’s modified Dulbecco’s media (IMDM) with 5% FBS, 50 U/ml penicillin, 50 μg/ml streptomycin, 2 mM L-glutamine, 0.1 mM non-essential amino acids, 1 mM sodium pyruvate and 50 μM 2-mercaptoethanol, and containing SCF (25 ng/ml), TPO (25 ng/ml) and Flt3-L (25 ng/ml), IL-11 (25 ng/ml), IL-3 (10 ng/ml), GM-CSF (10 ng/ml) and EPO (4 U/ml) as cytokines. To activate UPR, cells were treated with either tunicamycin (0.6 or 1.2 μg/ml; Sigma, T7765) for 16 hours or thapsigargin (0.05 or 0.1 μM; Sigma, T90330) for 6 hours and then plated onto the methylcellulose. For Xbp1 specific activation, cells were treated with 10 μM IXA4 (Selleck Chemicals, S9797) for 12 hours. For methylcellulose myeloid colony forming assays, 100 mouse cells were plated into methylcellulose (StemCell Technologies, M3231) as previously described^5^. Colonies were manually scored under a microscope after 6-8 days of culture. For human cells, 250 cells were cultured in StemSpan SFEM II media (StemCell Technologies, 09605) supplemented with StemSpan CD34+ expansion supplement (StemCell Technologies, 02691) and tunicamycin (0.6 μg/ml) for 12 hours and then plated onto methylcellulose (StemCell Technologies, M4034). Colonies were manually scored after 14 days of culture. For short-term *in vitro* differentiation of FcγR^−^ MPP3, cells were treated with 0.6 μg/ml tunicamycin for 6 or 12 hours and then cultured for extra 24 or 18 hours after washing out tunicamycin. For IXA4, cells were treated with 10 μM IXA4 for 24 hours before analysis.

### In vivo assays

For short-term *in vivo* differentiation assays, 20,000 FcγR^−^ MPP3 donor cells (CD45.2) isolated from littermate control and *Mx1-Cre::Jak2^V617F^* mice 12 weeks post-induction were retro-orbitally infused into recipient mice (CD45.1), which were analyzed for bone marrow (BM) contribution three days post infusion. Donor and recipient cells were identified by CD45.2-FITC and CD45.1-PE staining and recipient bone marrow cell population serves as an internal control for myeloid progenitor gating. For tunicamycin injection, mice were injected with tunicamycin (100 ng/g; IP) for five consecutive days before BM analysis. For repeated tunicamycin treatment, five consecutive day injection followed by five days off cycle was repeated. Tunicamycin was formulated in 1.5% DMSO (Sigma-Aldrich, D2650) and PBS. For ISRIB (MedChemExpress, HY-12495A) injection, mice were injected with ISRIB (1mg/kg; IP) for five consecutive days and then subjected to bleeding three days after last injection. ISRIB was formulated in 5% of DMSO, 60% of Kolliphor EL:ethanol (2:1 ratio), and 35% of saline. For transplantation experiments, recipient mice were lethally irradiated (10.5 Gy, delivered in split doses 4 hours apart) and injected retro-orbitally with 1 × 10^6^ BM cells.

### Immunofluorescence staining

For ATF4 staining, cells (2,000-3,000 cells/slide) were fixed, permeabilized, blocked and stained with a rabbit anti-mouse ATF4 (Cell Signaling, 11815S or Abcam, ab216839) primary antibody followed by a goat anti-rabbit-A488 (Invitrogen, A32731) secondary antibody. Slides were mounted with VectaShield Plus (Vector Laboratories, H-2000) containing 1 μg/ml DAPI.

### Bulk RNA-seq and bioinformatics analysis

RNA was isolated from 7,000 to 10,000 cells per sample using RNeasy Plus Micro Kit (Qiagen, 74034). RNA samples were submitted to the Genome Access Technology Center at the McDonnell Genome Institute for low input RNA sequencing. Differentially expressed gene (DEG) analysis was carried out using DEseq2^26^ package of R. Genes with an adjusted P-value less than 0.05 and fold change values greater than 1.5 were considered as DEGs. The enrichment of gene signatures based on KEGG pathway was examined using R package clusterProfiler^27^. RNA-seq data have been deposited in the Gene Expression Omnibus under accession code GSE297249.

## Results

### FcγR^−^ MPP3 is an erythroid-primed population

We previously showed that multipotent progenitor 3 (MPP3) is composed of two distinct subsets^5^. Myeloid-primed secretory ER^high^/FcγR^+^ MPP3 (hereafter FcγR^+^ MPP3) is a transitional population toward granulocyte/macrophage progenitor (GMP) commitment, while ER^low^/FcγR^−^ MPP3 (hereafter FcγR^−^ MPP3) represents the true multipotent part of the MPP3 compartment capable of generating FcγR^+^ MPP3 and differentiating towards other myeloid lineage fates like erythroid lineage^5^. We have shown that FcγR^+^ MPP3 are specifically expanded in *BCR-ABL* driven chronic myeloid leukemia (CML) mouse model, where granulocyte overproduction is a defining hallmark of the disease^5^. To determine whether MPP expansion was skewed in other forms of myeloproliferative neoplasm (MPN), we assessed various MPP sub-populations in a mouse model of polycythemia vera (PV) driven by the Jak2 mutant, *Jak2^V617F^*. Compared to controls, we found that both MPP2 and MPP3 populations were expanded in *Jak2^V617F^* mice and that this expansion of MPP3 was associated with a selective increase in FcγR^−^ MPP3, megakaryocyte/erythroid progenitor (MEP), and an overproduction of erythrocytes (Figure 1A; supplemental Figure 1A-C). These data, together with the specific expansion of FcγR^+^ MPP3 in CML mouse models^5^, show that MPP3 subsets are differentially expanded in different MPN mouse models based on the affected mature myeloid cells, mirroring the lineage potential of each MPP3 subset. To test whether Jak2 mutation augments the erythroid production potential of FcγR^−^ MPP3, we performed myeloid colony forming assays using FcγR^−^ MPP3 isolated from littermate control and *Jak2^V617F^* mice. FcγR^−^ MPP3 from *Jak2^V617F^* mice did not show enhanced erythroid colony forming capacity (Figure 1B). Furthermore, short-term *in vivo* differentiation assays showed that *Jak2^V617F^* FcγR^−^ MPP3 have a comparable level of MEP production (Figure 1C), indicating that *Jak2^V617F^* does not increase erythroid production of FcγR^−^ MPP3 per se. Rather, *Jak2^V617F^* increases the number of erythroid-primed FcγR^−^ MPP3, which results in erythroid overproduction. These data align with previous data showing that *BCR-ABL* increases granulocyte production not through enhancing intrinsic granulocyte potential of FcγR^+^ MPP3, but by expanding FcγR^+^ MPP3 population^5,6^. *Jak2^V617F^* also did not change the lineage potential of other MPP subsets (supplemental Figure 1D). In contrast, *Jak2^V617F^* HSCs had significantly increased erythroid colony formation, indicating that *Jak2^V617F^*mutation confers HSCs towards increased FcγR^−^ MPP3 and MPP2 numbers to promote erythroid production (Figure 1A; supplemental Figures 1A and D). To investigate whether *Jak2^V617F^* differentially affects HSCs and FcγR^−^ MPP3, we performed bulk RNA-sequencing (RNA-seq) using HSCs and FcγR^−^ MPP3 isolated from control and *Jak2^V617F^* mice. Principal component analysis (PCA) showed a clear segregation between control and *Jak2^V617F^* harboring cells (Figure 1D). Moreover, only 20% of the top 500 differentially expressed genes (DEGs) between control and *Jak2^V617F^* mice overlapped between HSCs and FcγR^−^ MPP3 (Supplemental Figure 1E). Indeed, KEGG pathway analysis on FcγR^−^ MPP3 DEGs uniquely showed alterations in metabolic pathway, indicating that *Jak2^V617F^* exerts distinct molecular effects in HSCs versus FcγR^−^ MPP3, though many cell cycle and DNA replication genes were commonly changed between both populations (Figure 1E; supplemental Figure 1F). Altogether, our data demonstrate that *Jak2^V617F^* drives HSCs to overproduce FcγR^−^ MPP3 and MPP2, which results in erythroid overproduction in a PV mouse model. These data also highlight the divided labor of MPP3 subsets in producing different myeloid lineage cells.

**Figure 1.**
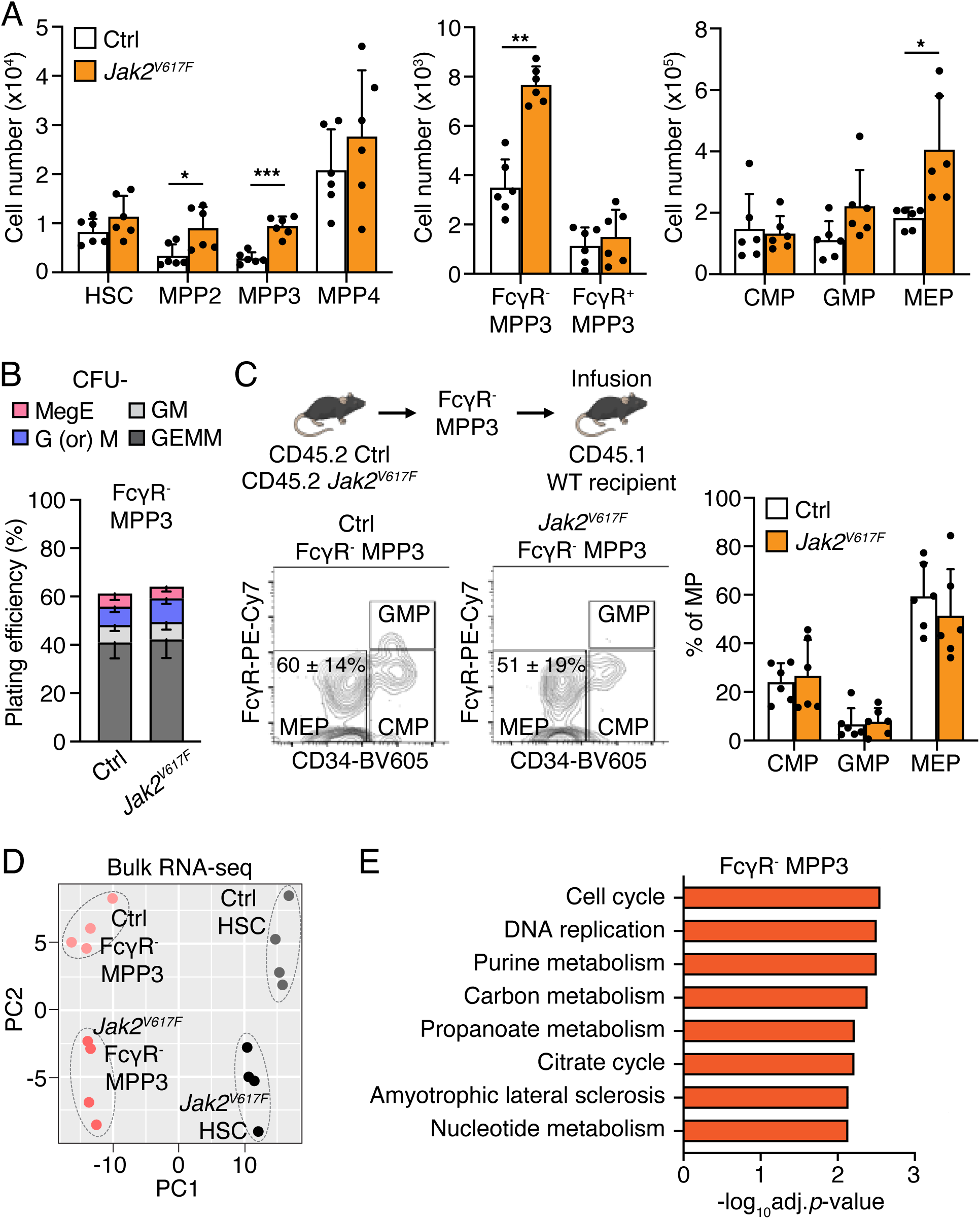
FcγR^−^ MPP3 expansion contributes to erythroid overproduction. (A) Hematopoietic stem and progenitor cell (HSPC) and myeloid progenitor (MP) population analyses of littermate control (Ctrl) and *Mx1-Cre::Jak2^V617F^* (*Jak2^V617F^*) mice 12 weeks post-induction. (B) Myeloid colony forming assays using FcγR^−^ MPP3 isolated from littermate control and *Mx1-Cre::Jak2^V617F^* mice 12 weeks post-induction (*n* = 8). Colony-forming units (CFUs) in methylcellulose assays were scored 8 days after plating. GEMM, granulocyte/erythroid/macrophage/megakaryocyte; GM, granulocyte/macrophage; G (or) M, granulocyte or macrophage; MegE, megakaryocyte/erythrocyte. (C) Short-term *in vivo* differentiation assays. Donor cells were isolated from littermate control and *Mx1-Cre::Jak2^V617F^* mice 12 weeks post induction (20,000 cells per mouse). Donor-derived cells were analyzed for MP (Lin^−^/Sca-1^−^/c-Kit^+^) contributions at 3 days post-infusion. Representative FACS plots and quantification of donor-derived frequencies are shown. (D) Principal component (PC) analysis of HSC and FcγR^−^ MPP3 bulk RNA-seq data from littermate control and *Mx1-Cre::Jak2^V617F^* mice. (E) KEGG pathway analysis of the top 500 differentially expressed genes (DEGs) in FcγR^−^ MPP3 from littermate control and *Mx1-Cre::Jak2^V617F^* mice. The DEG list is provided in Table S1. Data are means ± S.D., and statistical significance was assessed by a two-tailed unpaired Student’s *t-test*. * *p* ≤ 0.05, ** *p* ≤ 0.01, *** *p* ≤ 0.001.

### UPR signaling promotes myeloid cell production at the cost of erythroid lineage

To identify potential molecular regulators that dictate erythroid versus granulocyte/macrophage (G/M) output, we interrogated bulk RNA-seq data from FcγR^−^ MPP3 and FcγR^+^ MPP3^5^. From this analysis, we observed that several UPR-related genes (e.g., *Edem1, Calr, Hspa1a, Ero1l, Rpn1, Xbp1*) were enriched in FcγR^+^ MPP3, whereas erythroid genes (e.g., *Hbb-bt, Car2*) were enriched in FcγR^−^ MPP3. The full list of UPR-related genes in FcγR^+^ MPP3 and erythroid-related genes in FcγR^−^ MPP3 is provided in Table S3. XBP1, which is a key downstream mediator of the UPR, has been previously predicted to inhibit erythroid fate choice^28^. We therefore tested how chemical inducers of UPR activation influences the FcγR^−^ to FcγR^+^ MPP3 transition using myeloid colony forming assays. Treatment of wild type (WT) FcγR^−^ MPP3 with UPR inducers tunicamycin (Tuni) or thapsigargin (Thap) significantly increased granulocyte and macrophage colony formation while decreasing erythroid colony with overall reduced plating efficiency (Figures 2A and 2B). To make sure the decrease of erythroid colony is due to the transition of erythroid-primed FcγR^−^ MPP3 to G/M-biased FcγR^+^ MPP3, we performed apoptosis assays using both MPP3 subsets. UPR activator minimally impacted apoptosis in FcγR^−^ MPP3 with slightly higher level in FcγR^+^ MPP3, confirming that erythroid colony reduction upon UPR activation is not due to a selective killing of FcγR^−^ MPP3 (Supplemental Figure 2). UPR activators increased granulocyte and macrophage colony formation in FcγR^+^ MPP3 as well, indicating that UPR signaling also acts beyond FcγR^−^ to FcγR^+^ MPP3 transition to promote myeloid production (Figures 2C and 2D). To directly measure FcγR^+^ MPP3 production from FcγR^−^ MPP3, we conducted short-term *in vitro* differentiation assays as previously done^5^. Upon tunicamycin treatment, FcγR^+^ MPP3 production increased in a time dependent manner (Figure 2E).

**Figure 2.**
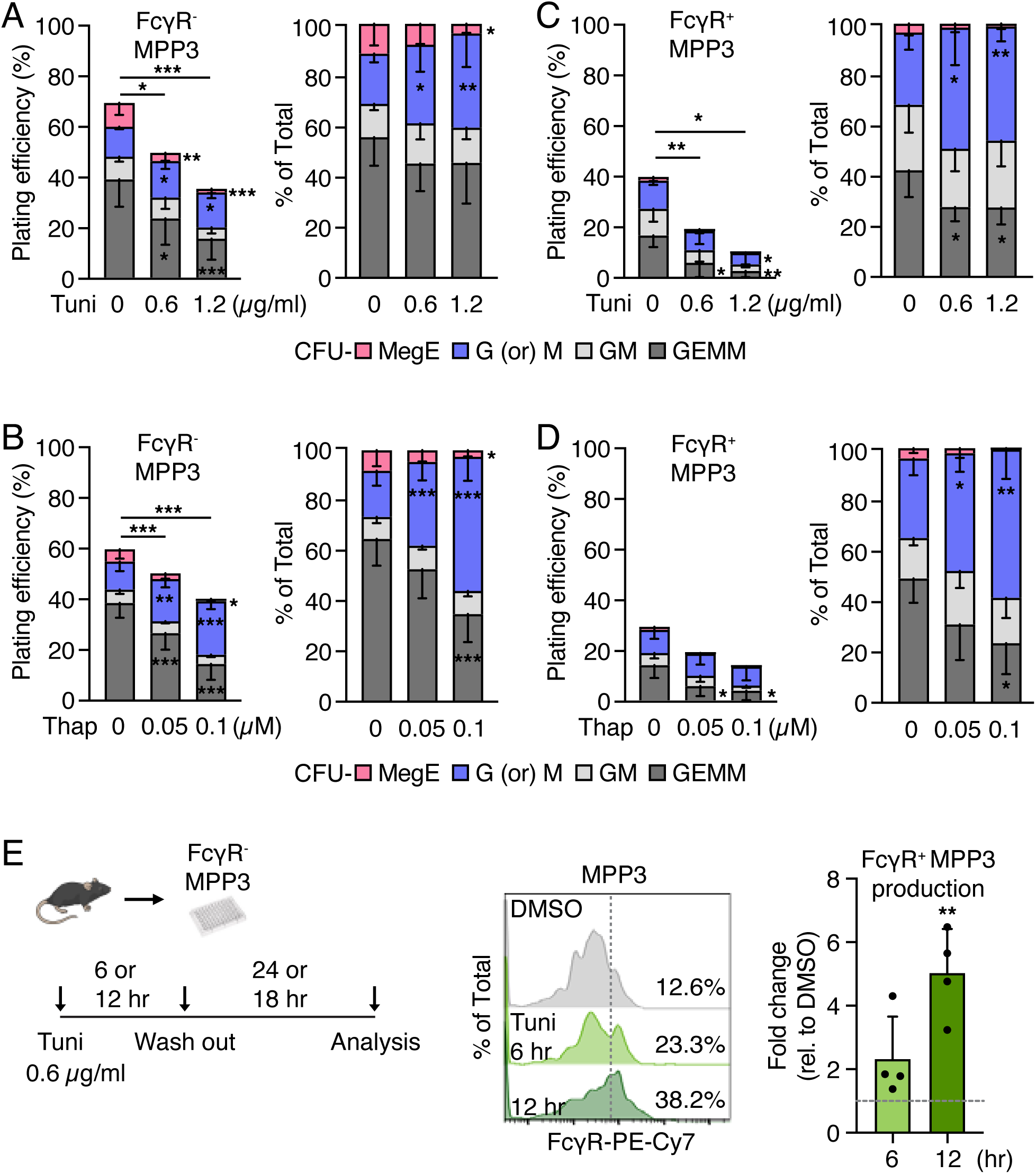
UPR signaling promotes FcγR^+^ MPP3 and myeloid cell production. (A and B) Myeloid colony forming assays using FcγR^−^ MPP3 treated with tunicamycin (Tuni) for 16 hours (*n* = 10) (A) or thapsigargin (Thap) for 6 hours (*n* = 8) (B). (C and D) Myeloid colony forming assays using FcγR^+^ MPP3 treated with Tuni for 16 hours (C) or Thap for 6 hours (D) (*n* = 5 for Tuni treatment and *n* = 4 for Thap treatment). Colony-forming units (CFUs) in methylcellulose assays were scored 6 days after plating. GEMM, granulocyte/erythroid/macrophage/megakaryocyte; GM, granulocyte/macrophage; G (or) M, granulocyte or macrophage; MegE, megakaryocyte/erythrocyte. Statistical significance is compared to control DMSO treatment. (E) Short-term *in vitro* differentiation of FcγR^−^ MPP3 upon tunicamycin (Tuni; 0.6 μg/ml) treatment. Cells (2,000 per well) were cultured for 6 or 12 hours (hr) with tunicamycin and cultured extra 24 or 18 hours, respectively, after washing out tunicamycin. Cells were then analyzed for HSPC markers. FACS plots and quantification of FcγR^+^ MPP3 production as a fold change relative to control (Ctrl: DMSO treated FcγR^−^ MPP3 cultured for total 30 hours) are shown. The gray dotted line indicates the fold change of 1. Data are means ± S.D., and statistical significance was assessed by a two-tailed unpaired Student’s *t-test*. * *p* ≤ 0.05, ** *p* ≤ 0.01, *** *p* ≤ 0.001.

To test whether chemical inducers of the UPR enhances non-pathogenic FcγR^+^ MPP3 production *in vivo*, we injected a low dose of tunicamycin (100 ng/g) for 5 days and analyzed bone marrow (BM) populations and peripheral blood (PB) (Figure 3A). The injection caused a slight decrease in body weight (4-5%) and reduced total MPP3, LSK (Lin^−^/Sca-1^+^/c-Kit^+^), and myeloid progenitor (MP) populations (Supplemental Figures 3A and 3B). However, within MPP3, FcγR^+^ MPP3 significantly increased (Figures 3B and 3C). GMP also significantly increased at the expense of MEP within MP followed by elevated PB myeloid cells (Figures 3D and 3E). We did not observe decreased erythrocytes in the PB right after tunicamycin injection, probably due to the longer lifespan of erythrocytes (45 to 50 days)^29,30^ and absolute high number. Therefore, we examined how multiple tunicamycin treatment cycles (five days on followed by five days off) impacted erythropoiesis (Figure 3F). While white blood cell and platelet numbers were unaffected by tunicamycin treatment, Hematocrit (HCT), the percentage of red blood cells (RBCs) in the blood, significantly decreased after one cycle and further decreased following additional treatments (Figure 3G; supplemental Figures 3C and 3D). Absolute RBC number also reduced upon tunicamycin treatment (Figure 3H). These results demonstrate that chemical inducers of the UPR facilitate FcγR^−^ to FcγR^+^ MPP3 conversion, which enhances myeloid cell production at the expense of erythroid cells.

**Figure 3.**
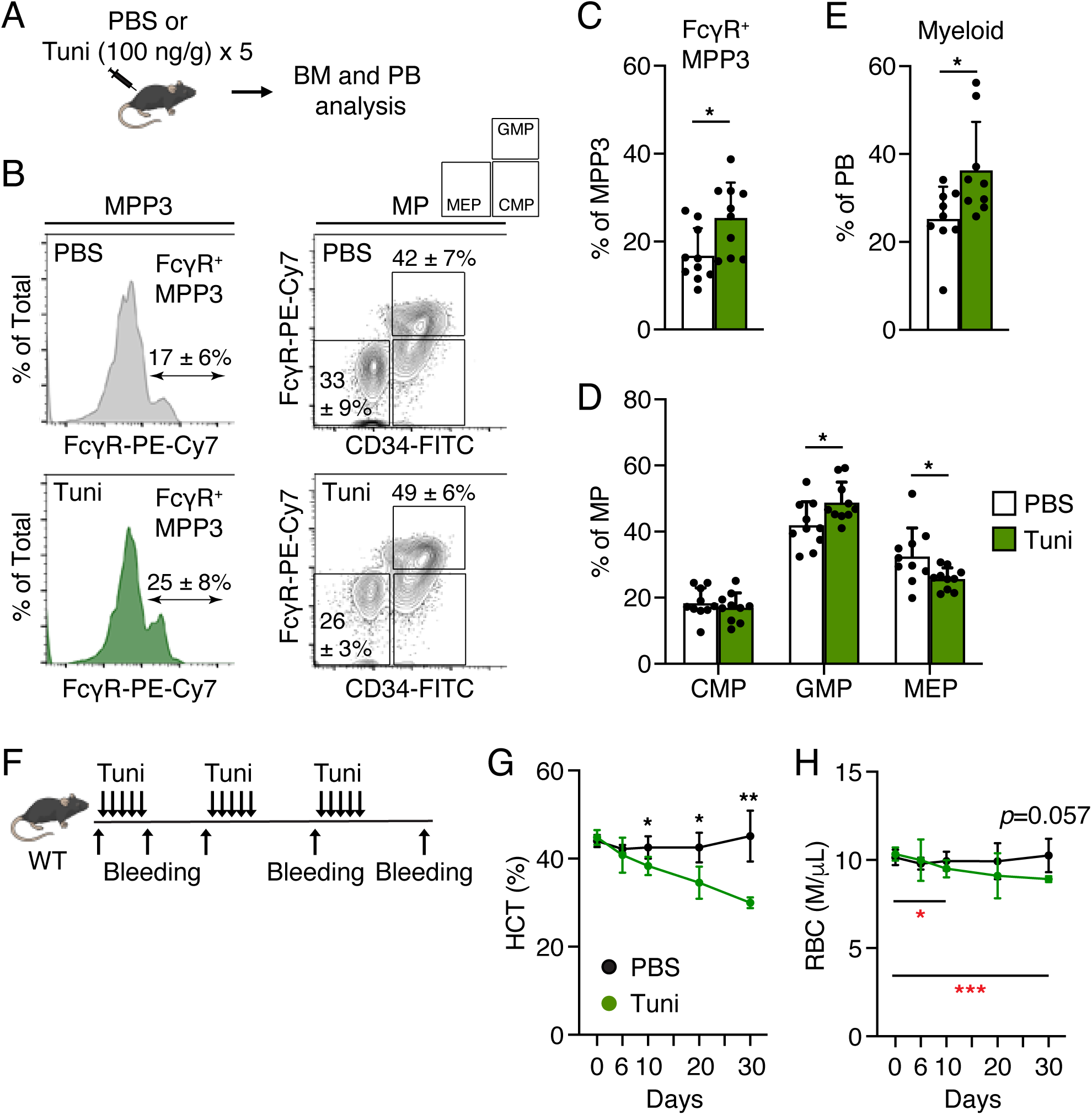
UPR signaling increases myeloid cell production at the expense of erythroid lineage. (A) Experimental scheme of *in vivo* tunicamycin (Tuni) injection. (B) Representative FACS plots showing the MPP3 and MP populations. (C-E) Quantification of FcγR^+^ MPP3 (C), MP population (D), and PB myeloid cells (E) upon tunicamycin treatment. Tuni, tunicamycin; BM, bone marrow; PB, peripheral blood; MP, myeloid progenitor. (F-H) Experimental scheme for repeated tunicamycin treatment in WT mice (F), peripheral blood (PB) hematocrit (HCT) (G) and red blood cell (RBC) (H) count upon repeated tunicamycin treatment (*n* = 4-6). Black asterisk denotes the significance between PBS and tunicamycin treatment. Red asterisk denotes the significance compared to day 0 within the Tuni treatment. Data are means ± S.D., and statistical significance was assessed by a two-tailed unpaired Student’s *t-test*. * *p* ≤ 0.05, ** *p* ≤ 0.01, *** *p* ≤ 0.001.

### UPR signaling cooperates with *Jak2^V617F^* mutation and increases erythroid production under a disease condition

To examine whether tunicamycin elicits similar effects under an erythroid-potentiating disease condition, we injected tunicamycin into primary *Jak2^V617F^*and littermate control mice. LSK and MP populations were significantly decreased in littermate controls but remained unchanged in *Jak2^V617F^* mice (Figure 4A). Although MPP3 proportion declined in both groups, FcγR^+^ MPP3 did not consistently increase in *Jak2^V617F^* mice (Figures 4B; supplemental Figures 4A and 4B). Instead, we observed variable responses by *Jak2^V617F^* mice upon tunicamycin injection, potentially reflecting differences in disease stage, cytokine signaling status, UPR sensitivity, or microenvironmental factors. Interestingly, MEP showed a significant increase at the cost of GMP followed by a higher RBC count in PB (Figures 4C and 4D). Of note, PB myeloid cells were still elevated, suggesting that UPR signaling also acted downstream of GMP as promoting myeloid output (Supplemental Figure 4C). This finding reveals that UPR signaling may function at multiple levels to enhance myelopoiesis. To corroborate these findings, we injected tunicamycin into transplanted *Jak2^V617F^* mice 6 weeks post-induction. Similar to the primary *Jak2^V617F^* mice, transplanted *Jak2^V617F^* mice showed heterogeneous responses and FcγR^+^ MPP3 did not invariably increase although LSK number reduced (Supplemental Figures 4D and 4E). MEP also increased with the loss of GMP, and PB myeloid cells were elevated, recapitulating the primary model (Supplemental Figures 4F and 4G).

**Figure 4.**
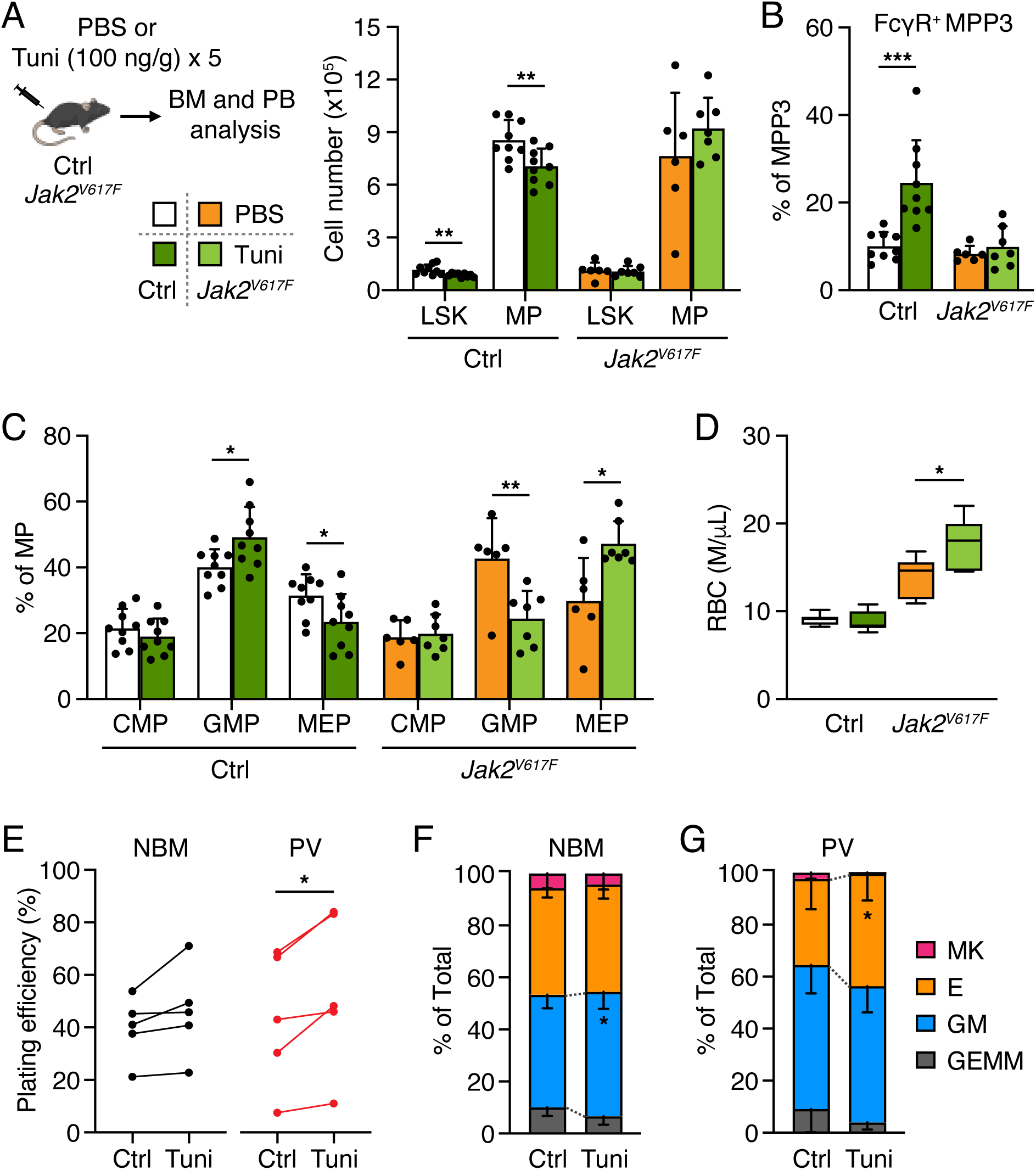
UPR signaling increases erythroid production in *Jak2^V617F^*mice. (A) Experimental scheme (left) and changes in LSK (Lin^−^/Sca-1^+^/c-Kit^+^) and myeloid progenitor (MP) populations in littermate control and *Jak2^V617F^* mice upon tunicamycin treatment (right). Data represent the combined results from both *Vav1-Cre* and *Mx1-Cre* driven *Jak2^V617F^*mice. (B-D) Quantification of FcγR^+^ MPP3 (B), MP population (C), and PB red blood cell (RBC) count (D) upon tunicamycin injection. Data are means + S.D., and statistical significance was assessed by a two-tailed unpaired Student’s *t-test*. (E-G) Overall plating efficiency (E) and myeloid colony forming capacity of normal bone marrow (NBM) (F) and PV patient bone marrow (PV) (G) MPP cells upon tunicamycin treatment (*n* = 5). Human bone marrow MPP cells were treated with tunicamycin (0.6 μg/ml) for 12 hours and then plated onto the methylcellulose for myeloid colony forming assays. Colonies were counted after 14 days of culture. GEMM, granulocyte/erythroid/macrophage/megakaryocyte; GM, granulocyte or macrophage; E, burst-forming unit-erythroid (BFU-E) and colony-forming unit-erythroid (CFU-E); MK, megakaryocyte. Data are means - S.D., and statistical significance was assessed by a two-tailed paired Student’s *t-test*. Tuni, tunicamycin; BM, bone marrow; PB, peripheral blood; Ctrl, control. * *p* ≤ 0.05, ** *p* ≤ 0.01, *** *p* ≤ 0.001.

To assess human relevance, we treated normal and PV patient BM MPPs (CD34^+^/CD38^−^/CD45RA^−^/CD90^−^/CD49f^−^)^31^ with tunicamycin (0.6 μg/ml for 12 hours)^12^ and performed myeloid colony forming assays. UPR activation increased GM colony formation in normal MPPs without reducing colony forming capacity (Figures 4E and 4F). PV patient MPPs, however, significantly increased erythroid output with higher colony forming capacity (Figures 4E and 4G). These data confirm that UPR signaling enhances myeloid output in normal hematopoiesis and reinforces the pathophysiology of *Jak2^V617F^* mutation in disease condition.

### UPR signaling promotes myelopoiesis through the XBP1 pathway

To identify the major UPR pathway that drives myeloid cell production and the potential mechanism underlying cooperative role of UPR signaling with *Jak2^V617F^* mutation, we focused on two downstream mediators of the UPR. We initially focused on XBP1 and ATF4 because our RNA-seq analysis revealed that *Xbp1* was enriched in G/M-committed FcγR^+^ MPP3, ATF6 induces *Xbp1* mRNA, and hematopoietic ATF4 deficient mice have erythroid defects ^5,21,23,32^. We treated WT MPP3 cells with tunicamycin (1.2 μg/ml for 12 hr) or thapsigargin (0.05 μM for 6 hr) and measured spliced and unspliced *Xbp1* transcripts using qRT-PCR. Increased ratio of spliced *Xbp1* over unspliced form is regarded as an indication of XBP1 transcriptional activity^32^. The ratio of spliced *Xbp1* was increased in both tunicamycin and thapsigargin-treated MPP3 (Figure 5A). Importantly, tunicamycin treatment also induced *Xbp1* transcription with increased spliced *Xbp1* transcripts, which produce XBP1 transcription factor that moves to the nucleus^32^ (Figure 5A). For the ATF4 pathway, we performed immunofluorescence assays and quantified nuclear ATF4 levels. Tunicamycin treatment activated ATF4 pathway, while thapsigargin treatment did not in MPP3 cells (Figure 5B). These data showed that only XBP1 pathway was activated by both tunicamycin and thapsigargin treatment in MPP3 cells, indicating that XBP1 is involved in the G/M priming of MPP3 cells. We then directly tested whether increasing XBP1 activity promotes myeloid cell production by performing myeloid colony forming assays using IXA4, a highly selective, non-toxic chemical that stimulates IRE1α, which is the upstream activator of XBP1s^33^. Upon IXA4 treatment (10 μM for 12 hr), MPP3 significantly increased G/M colony formation, similar to thapsigargin treatment but without compromising overall colony forming capacity (Figure 5C). Short-term *in vitro* differentiation assays also confirmed that IXA4 treatment significantly increased FcγR^+^ MPP3 production from FcγR^−^ MPP3 (Figure 5D). We also used *Xbp1* conditional knockout mice (*Xbp1* KO; *Mx1-Cre::Xbp1^fl/fl^*) and performed myeloid colony forming assays. FcγR^−^ MPP3 isolated from littermate controls and 8 weeks post-induction *Xbp1* KO were treated with thapsigargin (0.05 μM for 6 hours). Thapsigargin treatment significantly increased G or M colonies while reducing erythroid colonies in controls as repeatedly shown in this study (Figure 5E). However, *Xbp1* deficient FcγR^−^ MPP3 did not produce more G or M colonies upon UPR activation (Figure 5E). Of note, *Xbp1* deficient FcγR^−^ MPP3 formed significantly higher number of erythroid colonies compared to controls and did not reduce erythroid colony forming capacity upon thapsigargin treatment (Figure 5E). This data aligns with the previous study predicting *Xbp1* to inhibit erythroid fate^28^ and reinforces the idea that *Xbp1* acts as a regulator of myeloid versus erythroid fate decision in FcγR^−^ MPP3 by promoting FcγR^−^ MPP3 transition to FcγR^+^ MPP3 (Figure 5F). Taken together, these data demonstrate that UPR signaling promotes myelopoiesis through the XBP1 pathway.

**Figure 5.**
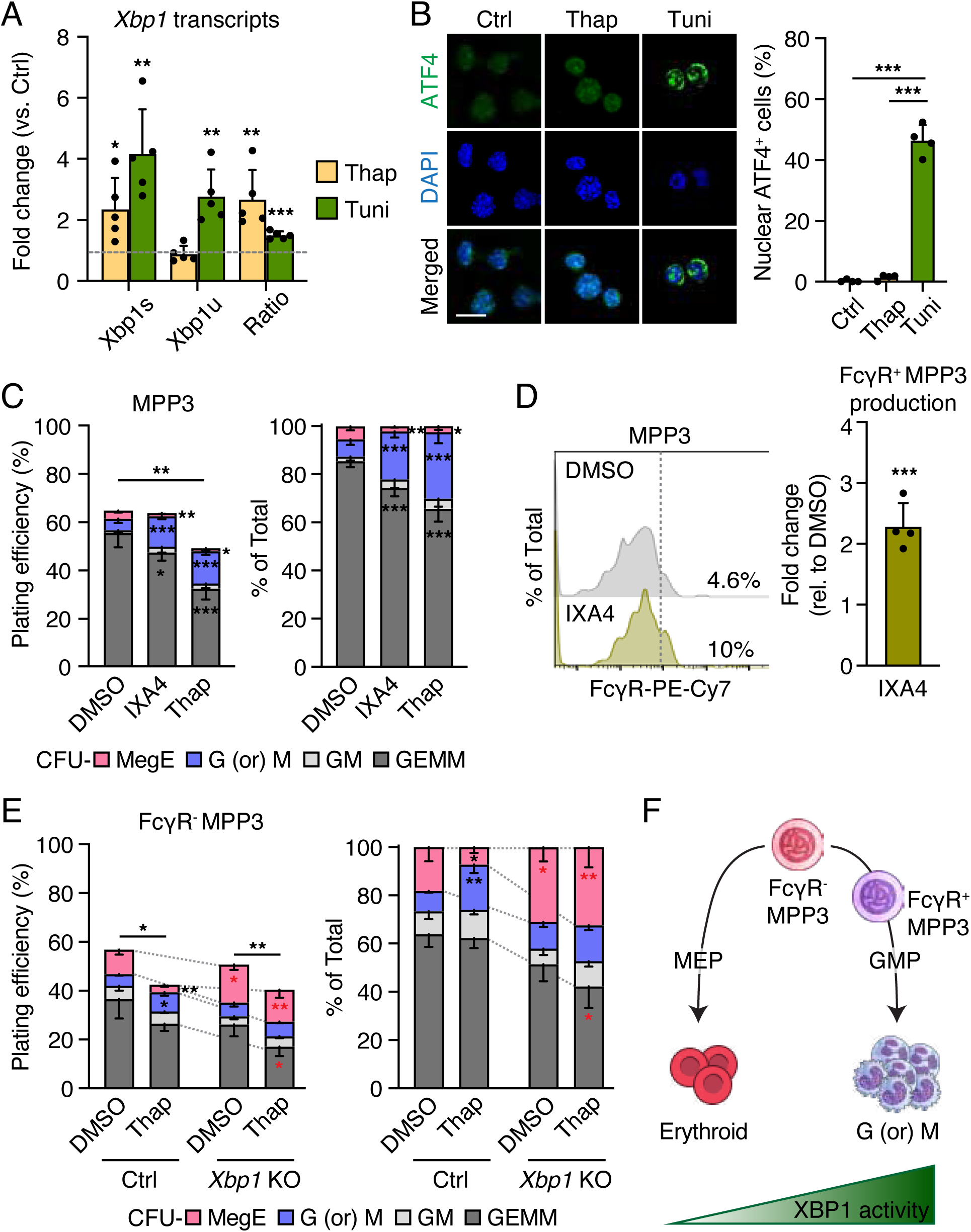
UPR signaling promotes myelopoiesis through the XBP1 pathway. (A) Quantitative RT-PCR analyses of *Xbp1* transcripts of MPP3 upon tunicamycin (Tuni) (1.2 μg/ml for 12 hr) or thapsigargin (Thap) (0.05 μM for 6 hr) treatment. DMSO treated MPP3 cells served as controls. Fold change relative to control samples are shown. Dotted line denotes the fold change of 1. Statistical significance is compared to controls. Xbp1s: spliced *Xbp1* transcript; Xbp1u: unspliced *Xbp1* transcript; Ratio: ratio of spliced over unspliced *Xbp1* transcript. (B) Representative images and quantification of nuclear ATF4 positive MPP3 cells upon tunicamycin (1.2 μg/ml for 12 hr) or thapsigargin (0.05 μM for 6 hr) treatment. DMSO treated MPP3 cells served as controls. Scale bar: 10 μm. (C) Quantification of myeloid colony formation of MPP3 cells upon IXA4 (10 μM) and thapsigargin (Thap; 0.05 μM) treatment for 12 hours (*n* = 4). CFU, colony forming unit; MegE, megakaryocyte/erythrocyte; G (or) M, granulocyte or macrophage; GM; granulocyte/macrophage, GEMM, granulocyte/erythroid/macrophage/megakaryocyte. (D) Short-term *in vitro* differentiation of FcγR^−^ MPP3 upon IXA4 treatment. Cells (2,000 per well) were cultured with IXA4 (10 μM) for 24 hours and then analyzed for HSPC markers. Representative FACS plots and quantification of FcγR^+^ MPP3 production as fold change relative to control (Ctrl: DMSO treated FcγR^−^ MPP3) are shown. Statistical significance is compared to controls. (E) Myeloid colony forming assays using FcγR^−^ MPP3 isolated from littermate control (Ctrl) and *Xbp1* conditional knockout mice (*Xbp1* KO) treated with thapsigargin (Thap; 0.05 μM) for 6 hours (*n* = 3). Black asterisk denotes the significance between DMSO and thapsigargin treatment within the same cell type. Red asterisk denotes the significance between Ctrl and *Xbp1* KO cells within the same treatment. (F) Model depicting the function of XBP1 as a regulator of myeloid versus erythroid fate choice in FcγR^−^ MPP3 by promoting FcγR^−^ MPP3 transition to FcγR^+^ MPP3. G (or) M, granulocyte or macrophage. Model is made with NIH BioArt. Data are means ± S.D., and statistical significance was assessed by a two-tailed unpaired Student’s t-test. * *p* ≤ 0.05; ** *p* ≤ 0.01; *** *p* ≤ 0.001.

### ATF4 collaborates with *Jak2^V617F^* and enhances erythropoiesis

To investigate ATF4 activity in *Jak2^V617F^* mice upon chemical activation of the UPR, we performed ATF4 immunofluorescence assays using MPP3 cells isolated from vehicle versus tunicamycin treated littermate control and *Jak2^V617F^*mice (Figure 6A). Tunicamycin (100 ng/g) was injected for five consecutive days before MPP3 isolation. Similar to *in vitro* tunicamycin treatment, MPP3 cells from tunicamycin injected mice had significantly higher nuclear ATF4 levels in controls (Figure 6A). In *Jak2^V617F^* mice, nuclear ATF4 level was already significantly higher compared to littermate controls (Figure 6A). This data indicates that *Jak2^V617F^* activates ATF4 pathway, which likely contributes to erythroid overproduction given that ATF4 promotes erythropoiesis through ribosome biogenesis^21^. Our molecular data also reflects this ATF4 activation in MPP3 cells (e.g., purine metabolism, carbon metabolism, citrate cycle, nucleotide metabolism) as ATF4 activates metabolic pathway genes such as genes in amino acid metabolism, nucleotide synthesis and carbon metabolism^34^ (Figure 1E). Tunicamycin injection further increased ATF4 activity in *Jak2^V617F^* mice, suggesting that ATF4 hyperactivation upon tunicamycin injection help *Jak2^V617F^* even more overproduce erythroid cells (Figures 4D and 6A). Of note, littermate control mice with tunicamycin injection also had high nuclear ATF4 level but they did not increase RBC production, indicating that ATF4 activation is not sufficient to promote erythropoiesis (Figures 4D and 6A). To directly test whether ATF4 activation in *Jak2^V617F^* mice promotes erythropoiesis, we treated transplanted *Jak2^V617F^* mice with ISRIB, an eIF2 modifier that results in ATF4 inhibition^35,36^ (1mg/kg) for five days and measured RBC three days post injection (Figure 6B). ISRIB treated *Jak2^V617F^*mice significantly decreased RBC number while maintaining other blood counts (Figures 6B-D). In primary *Jak2^V617F^* mice, ISRIB injection also reduced RBC, but littermate control mice injected with ISRIB showed no difference in RBC number, indicating that lowering ATF4 activity can be used to ameliorate erythroid overproduction in *Jak2^V617F^*driven PV (Figure 6E). Altogether, these results show that UPR signaling enhances erythroid production in collaboration with *Jak2^V617F^* via the ATF4 pathway (Figure 6F).

**Figure 6.**
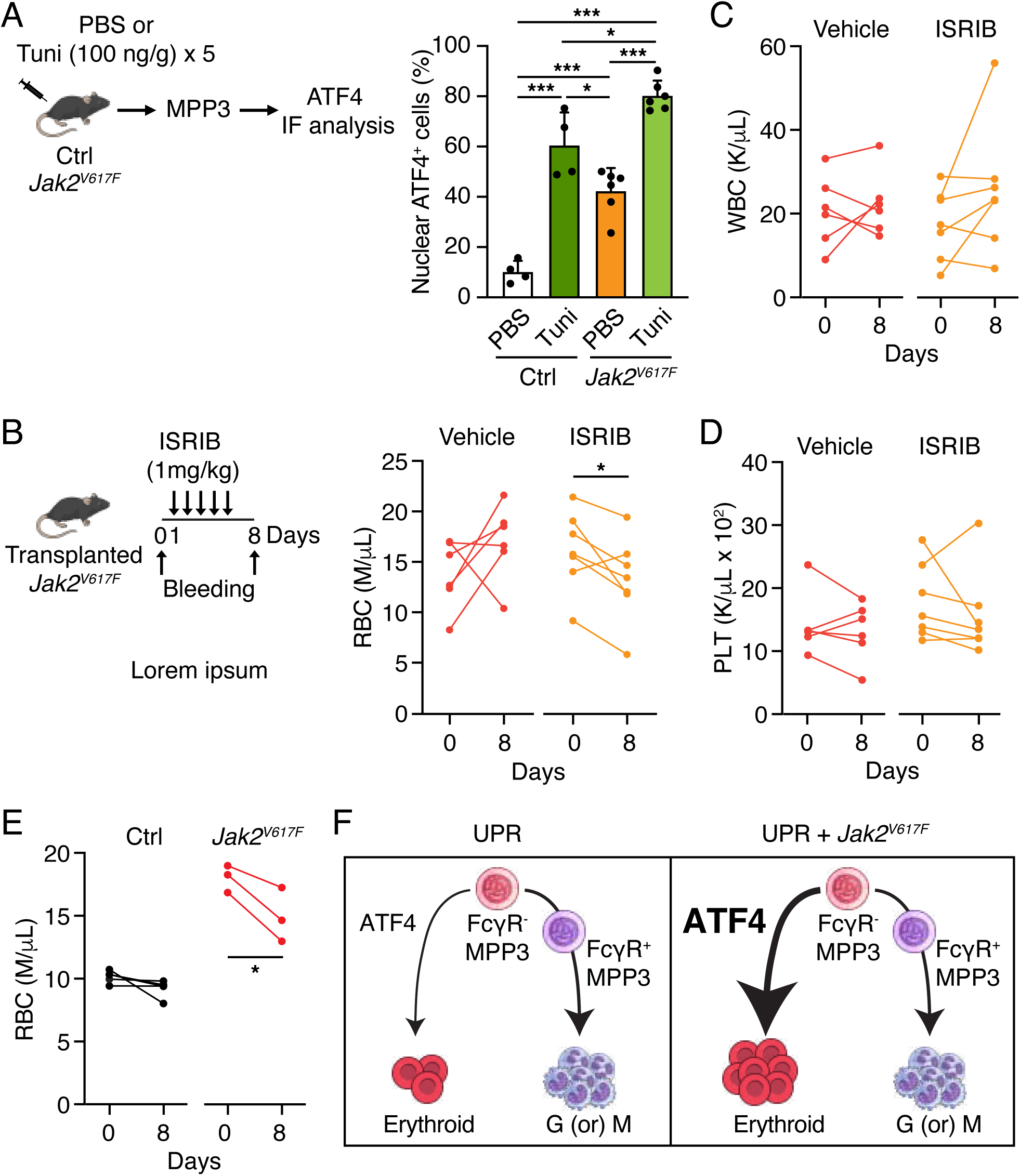
UPR signaling cooperates with *Jak2^V617F^* and increases erythroid production via the ATF4 pathway. (A) Experimental scheme and quantification of nuclear ATF4 positive MPP3 cells isolated from vehicle versus tunicamycin (Tuni) treated control (Ctrl) and *Jak2^V617F^*mice. Data are means + S.D., and statistical significance was assessed by a two-tailed unpaired Student’s *t-test*. (B-D) Experimental scheme and red blood cell (RBC) (B), white blood cell (WBC) (C) and platelet (PLT) (D) count upon ISRIB injection in transplanted *Jak2^V617F^*mice. Transplanted *Jak2^V617F^* mice were prepared as shown in supplemental Figure 4D. Mice were injected with ISRIB (1mg/kg) 6 weeks post-induction. Statistical significance was assessed by a two-tailed paired Student’s *t-test*. (E) RBC count upon ISRIB injection (1mg/kg) in littermate control and *Jak2^V617F^* mice. Both groups were injected with ISRIB. Injection scheme is shown in Figure 6B. Ctrl, littermate control mice. Statistical significance was assessed by a two-tailed paired Student’s *t-test*. (F) Model depicting the collaborative work of UPR signaling and *Jak2^V617F^* mutation in overproducing erythroid cells by hyperactivating ATF4. G (or) M, granulocyte or macrophage. Model is made with NIH BioArt. * *p* ≤ 0.05; *** *p* ≤ 0.001.

## Discussion

Recent studies demonstrated that UPR signaling controls HSPC biology^12–14,20–22^. Whether UPR signaling regulates lineage fate decision in the hematopoietic system remains unclear. Considering UPR is elevated in many cancers and targeting UPR signaling is considered as therapeutic interventions^11^, whether UPR signaling alters hematopoietic lineage output is an important consideration when targeting UPR in cancers. Here, we found that chemical activation of the UPR promotes myelopoiesis by increasing the transition of erythroid-primed FcγR^−^ MPP3 to myeloid-committed FcγR^+^ MPP3, which leads to GMP expansion followed by myeloid cell increase in PB through the XBP1 pathway. These increased myeloid cells come at the cost of erythroid production, indicating that MPP3 is the divergent point of myeloid versus erythroid lineage specification. These data also suggest that UPR signaling controls myeloid versus erythroid output in the hematopoietic system. Especially, sustained UPR activation progressively decreased RBC level in the body, implying that prolonged up-regulation of UPR signaling itself can cause anemia in hematological malignancies.

In a mouse model of PV, UPR signaling cooperates with *Jak2^V617F^* mutation and increases erythroid cells via the ATF4 pathway. ATF4 is already activated in *Jak2^V617F^* mice as evidenced by up-regulation of canonical ATF4 metabolic target genes (e.g., *Phgdh*, *Shmt1*, *Ass1*, *Psat1*, etc.) in *Jak2^V617F^* FcγR^−^ MPP3 and high nuclear ATF4 level in *Jak2^V617F^* MPP3. Consistent with this observation, Jak2 signaling has been shown to activate protein kinase R (PKR), a key eIF2α kinase, which, in turn, can activate ATF4^37,38^. We show that ATF4 exerts its effect on erythroid-primed FcγR^−^ MPP3 and ATF4 hyperactivation further increases erythroid overproduction in *Jak2^V617F^* mice. Of note, high ATF4 activity induced by UPR activation alone did not increase RBC number, reinforcing the idea that ATF4 activity itself is not sufficient although it is necessary to increase erythropoiesis in collaboration with *Jak2^V617F^*. How much ATF4 activity contributes to *Jak2^V617F^* other phenotypes will be of interest to consider targeting ATF4 as a therapeutic intervention in PV.

Upon tunicamycin injection, some *Jak2^V617F^* mice increased FcγR^+^ MPP3 but some *Jak2^V617F^* mice did not. This could be possibly because the tunicamycin dose we injected was not strong enough to overpower the underlying driving force of the *Jak2^V617F^* mutation on erythroid lineage. Although GMP was relatively reduced in the MP compartment, the PB myeloid cells still significantly increased, indicating that myelopoiesis is not just a linear cascading process and that UPR signaling may also act on downstream of GMP to enhance myelopoiesis potentially through its canonical pro-survival function. This also aligns with our *in vitro* myeloid colony forming assay data with FcγR^+^ MPP3, which showed increased G/M colony formation upon UPR activation. The increase of PB myeloid cells also implies that *Jak2^V617F^* may provide a self-enforcing feedback loop, which causes the progression of PV to more advanced diseases like myelofibrosis by constantly activating UPR signaling. Recent study revealed that type 2 calreticulin mutations activate ATF6 to enhance survival in MPN^39^, providing another evidence that disease-causing mutation collaborates with UPR signaling for pathogenesis. Further investigation into whether UPR components interact with other hematologic/oncogenic mutations and their functional consequences will provide valuable therapeutic insights. Taken together, our study uncovers that distinct UPR pathways function in lineage-specific production in a context-dependent manner, wherein UPR signaling promotes myelopoiesis through the XBP1 pathway under normal conditions, while enhancing erythropoiesis via the ATF4 pathway in collaboration with *Jak2^V617F^* mutation.

## Supporting information

Combined supplemental tables

## Acknowledgements

We thank Dr. Link (Wash U) for *UBC-Cre* mice, Dr. Glimcher (Dana-Farber Cancer Institute) for *Xbp1^fl/fl^* mice, and Lina Joo (Wash U) for technical assistance. This work was supported by NIH grant K01 DK120780, R56DK139218, Leukemia Research Foundation P22-05724, American Cancer Society IRG-21-133-64-01 and CAT-24-1377095-01-CAT, NCI Specialized Program of Research Excellence in Leukemia (SPORE) DRP-2203, the Alvin J. Siteman Cancer Center through The Foundation for Barnes-Jewish Hospital AW00014358 to Y-A. Kang and NIH grant R01HL134952 to S.T. Oh.

## Authorship and conflict-of-interest statements

Contribution: H. Choi and Y-A. Kang designed the study; H. Choi, S.E. Jung, Y. Earth, J. Yi and Y-A. Kang performed experiments, collected and analyzed data; H. Paik performed all the bioinformatic analyses; M.J. Cox and S.T. Oh provided PV patient samples; S.T. Oh provided *Vav1-Cre::Jak2^V617F^* mice; S.M. Sykes provided reagents and critical advice for experimental design; H. Choi and Y-A. Kang wrote the manuscript; H. Choi, S.E. Jung, S.M. Sykes, S.T. Oh and Y-A. Kang edited the manuscript. All authors read and approved the manuscript.

Disclosure of Conflicts of Interest: H. Choi, S.E. Jung, Y. Earth, J. Yi, M. J. Cox, H. Paik, S.M. Sykes, Y-A. Kang declare no competing financial interests. S.T. Oh has served as a consultant for Kartos Therapeutics, CTI BioPharma, Celgene–Bristol Myers Squibb, Disc Medicine, Blueprint Medicines, PharmaEssentia, Constellation, Geron, AbbVie, Sierra Oncology and Incyte. S.E. Jung’s current affiliation: Seoul National University, Seoul, South Korea.

## Data Sharing Statement

Data are available upon request from the corresponding author, Yoon-A Kang. RNA-seq data have been deposited in the Gene Expression Omnibus under accession code GSE297249.

## Tables

Supplementary Table 1. Top 500 differentially expressed gene (DEG) list between control and *Jak2^V617F^*FcγR^−^ MPP3

Supplementary Table 2. Top 500 DEG list between control and *Jak2^V617F^* HSCs

Supplementary Table 3. List of UPR-related genes in FcγR^+^ MPP3 and erythroid-related genes in FcγR^−^ MPP3 from bulk RNA-seq data between FcγR^−^ MPP3 and FcγR^+^ MPP3

Supplementary Table 4. Patient sample information

Supplementary Table 5. List of antibodies used

---

**Figure S1.**
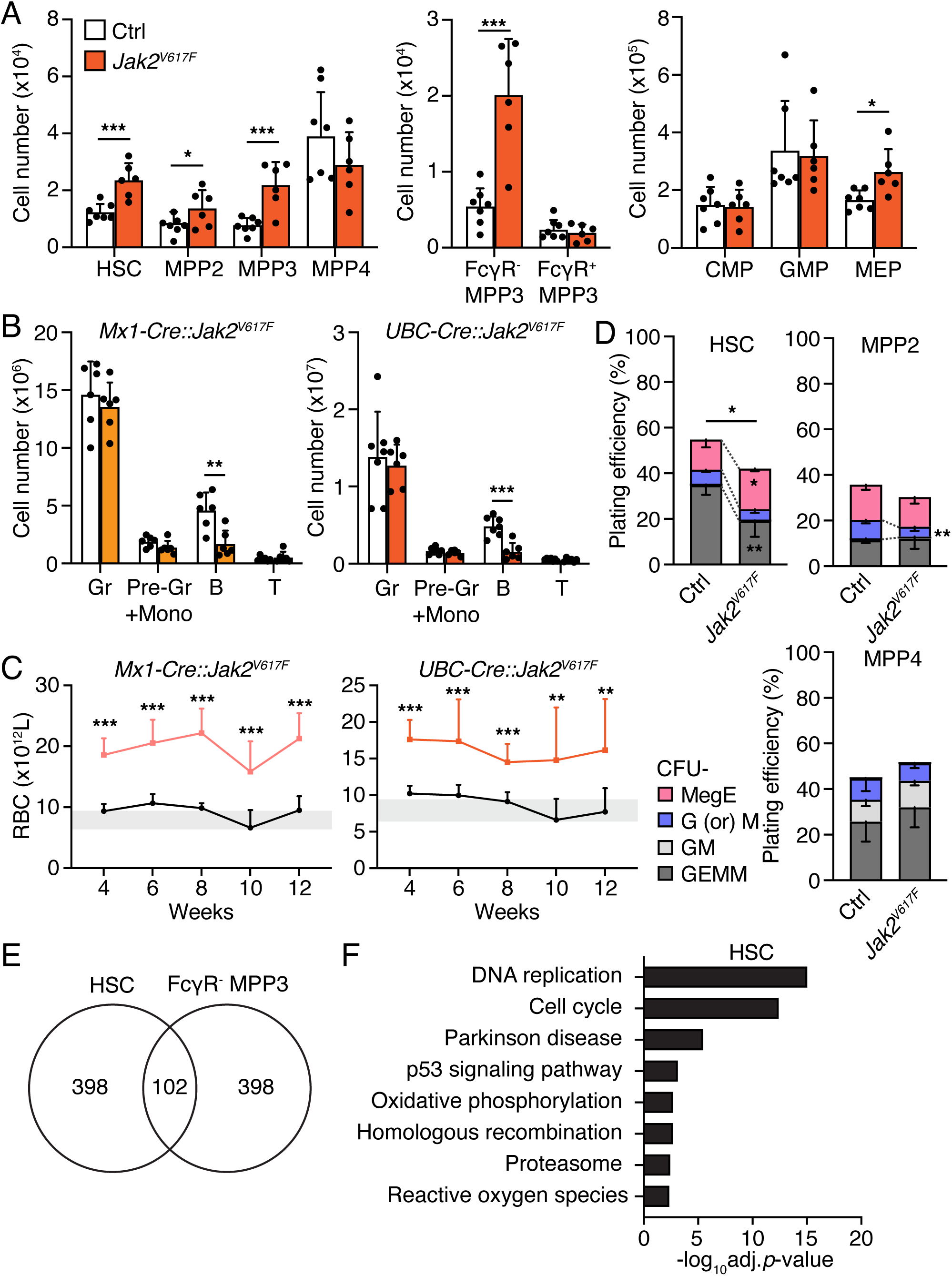
*Jak2^V617F^*mutation differentially affects HSCs and FcγR^−^ MPP3 population. (A) Hematopoietic stem and progenitor cell (HSPC) and myeloid progenitor (MP) population analyses of littermate control (Ctrl) and *UBC-Cre::Jak2^V617F^*(*Jak2^V617F^*) mice 12 weeks post-induction. (B) Mature bone marrow population analysis of littermate controls and *Mx1-Cre::Jak2^V617F^*mice (left) or *UBC-Cre::Jak2^V617F^* mice (right). Gr, granulocytes; Pre-Gr + Mono, pre-granulocytes and monocytes; B, B cells; T, T cells. (C) Red blood cell (RBC) counts upon *Jak2^V617F^* induction in *Mx1-Cre::Jak2^V617F^* mice (left; *n* = 9 for control and 7 for *Jak2^V617F^* up to 10 weeks and *n* = 19 for control and 22 for *Jak2^V617F^* at 12 weeks) and *UBC-Cre::Jak2^V617F^* mice (right; *n* = 13 for control and 6 for *Jak2^V617F^*). (D) Myeloid colony forming assays using HSC, MPP2, and MPP4 isolated from littermate control and *Mx1-Cre::Jak2^V617F^* mice 12 weeks post-induction (*n* = 5). Colony-forming units (CFUs) in methylcellulose assays were scored 8 days after plating. GEMM, granulocyte/erythroid/macrophage/megakaryocyte; GM, granulocyte/macrophage; G (or) M, granulocyte or macrophage; MegE, megakaryocyte/erythrocyte. (E) Venn diagram showing the overlap of differentially expressed genes (DEGs) between HSC and FcγR^−^ MPP3 from the top 500 DEG list between control and *Jak2^V617F^* mice in each cell type. 500 DEG from control HSC versus *Jak2^V617F^* HSC was compared with 500 DEG from control FcγR^−^ MPP3 versus *Jak2^V617F^* FcγR^−^ MPP3. (F) KEGG pathway analysis of DEGs in HSCs between littermate control and *Mx1-Cre::Jak2^V617F^*bulk RNA-seq data. The top 500 DEG list is presented in Table S2. Data are means ± S.D., and statistical significance was assessed by a two-tailed unpaired Student’s *t-test*. * *p* ≤ 0.05, ** *p* ≤ 0.01, *** *p* ≤ 0.001.

**Figure S2.**
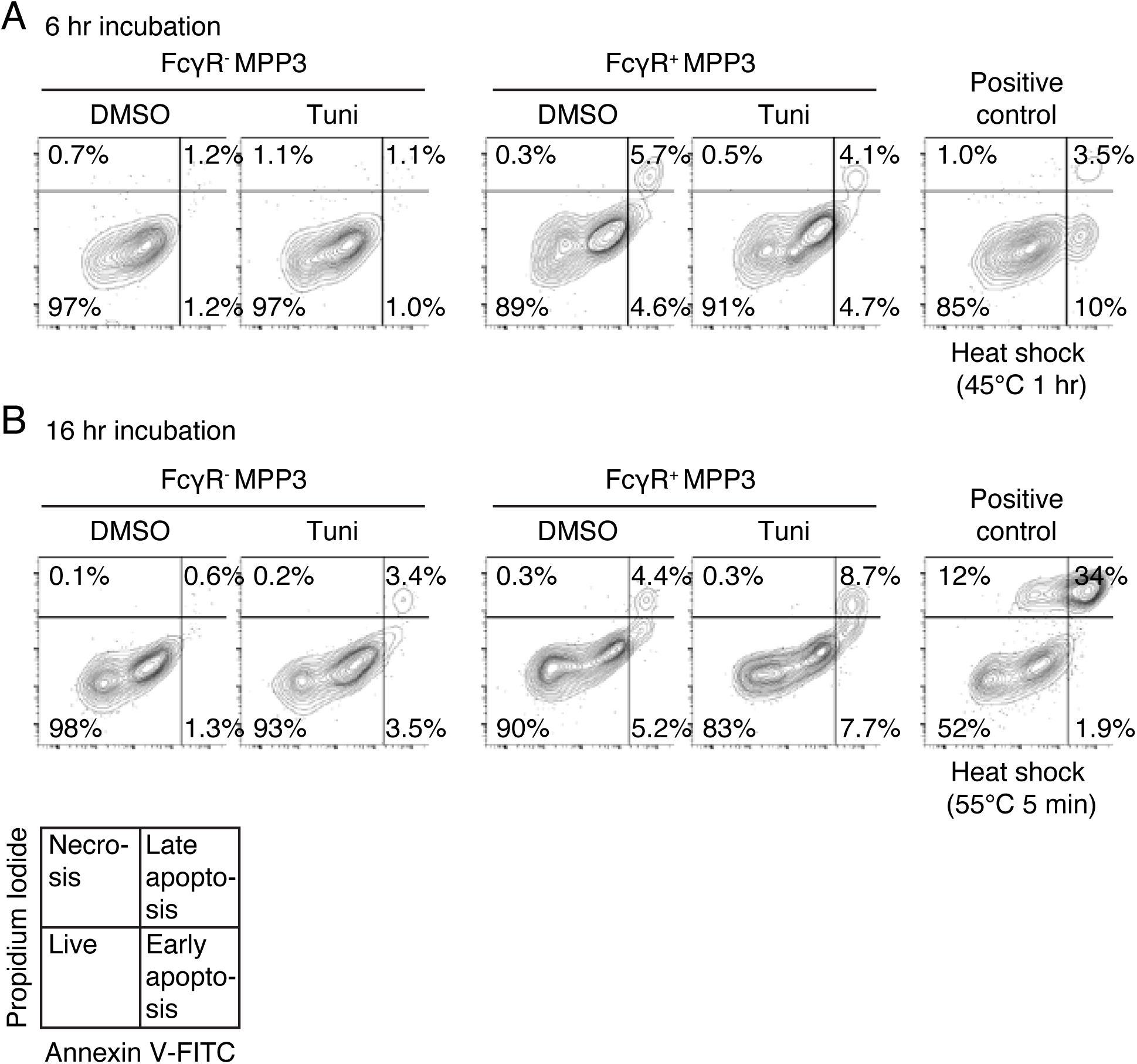
UPR signaling does not selectively cause apoptosis in FcγR^−^ MPP3. Representative FACS plots showing the apoptotic cells upon tunicamycin (Tuni) treatment (0.6 μg/ml) in MPP3 subsets for 6 hours (A) and 16 hours (B) (*n* = 2). Cultured FcγR^−^ MPP3 with heat shock serves as positive controls. hr, hours; min, minutes.

**Figure S3.**
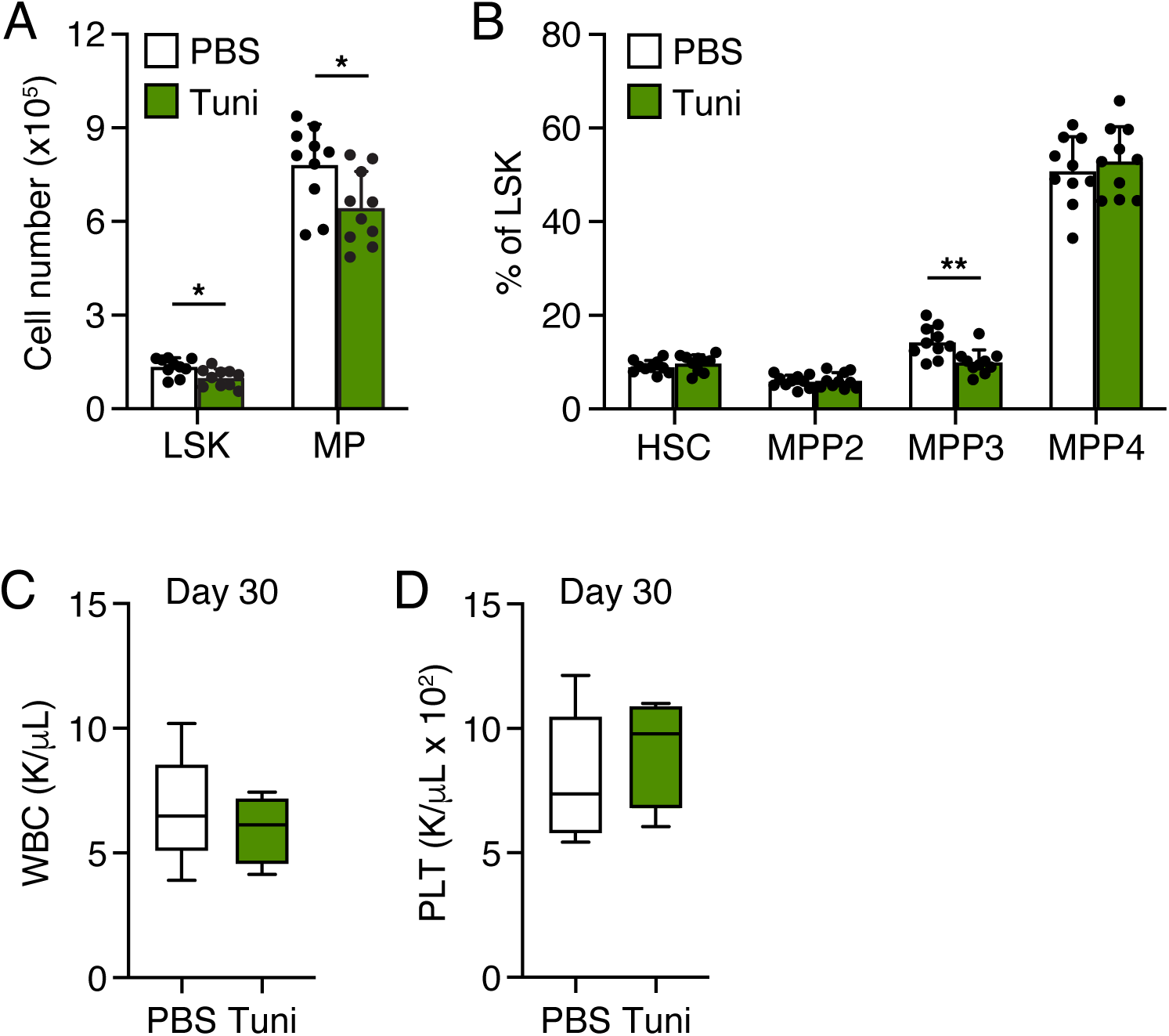
UPR activation changes bone marrow population. (A) Changes in LSK (Lin^−^/Sca-1^+^/c-Kit^+^) and myeloid progenitor (MP) populations in wild type (WT) mice upon tunicamycin (Tuni) treatment. (B) Quantification of hematopoietic stem and progenitor cell (HSPC) populations in wild type mice upon tunicamycin treatment. Results are expressed as a percentage of LSK. (C and D) Peripheral blood (PB) white blood cell (WBC) (C) and platelet (PLT) (D) counts upon repeated tunicamycin treatment in WT mice (*n* = 4-6). Data are means ± S.D., and statistical significance was assessed by a two-tailed unpaired Student’s *t-test*. * *p* ≤ 0.05, ** *p* ≤ 0.01.

**Figure S4.**
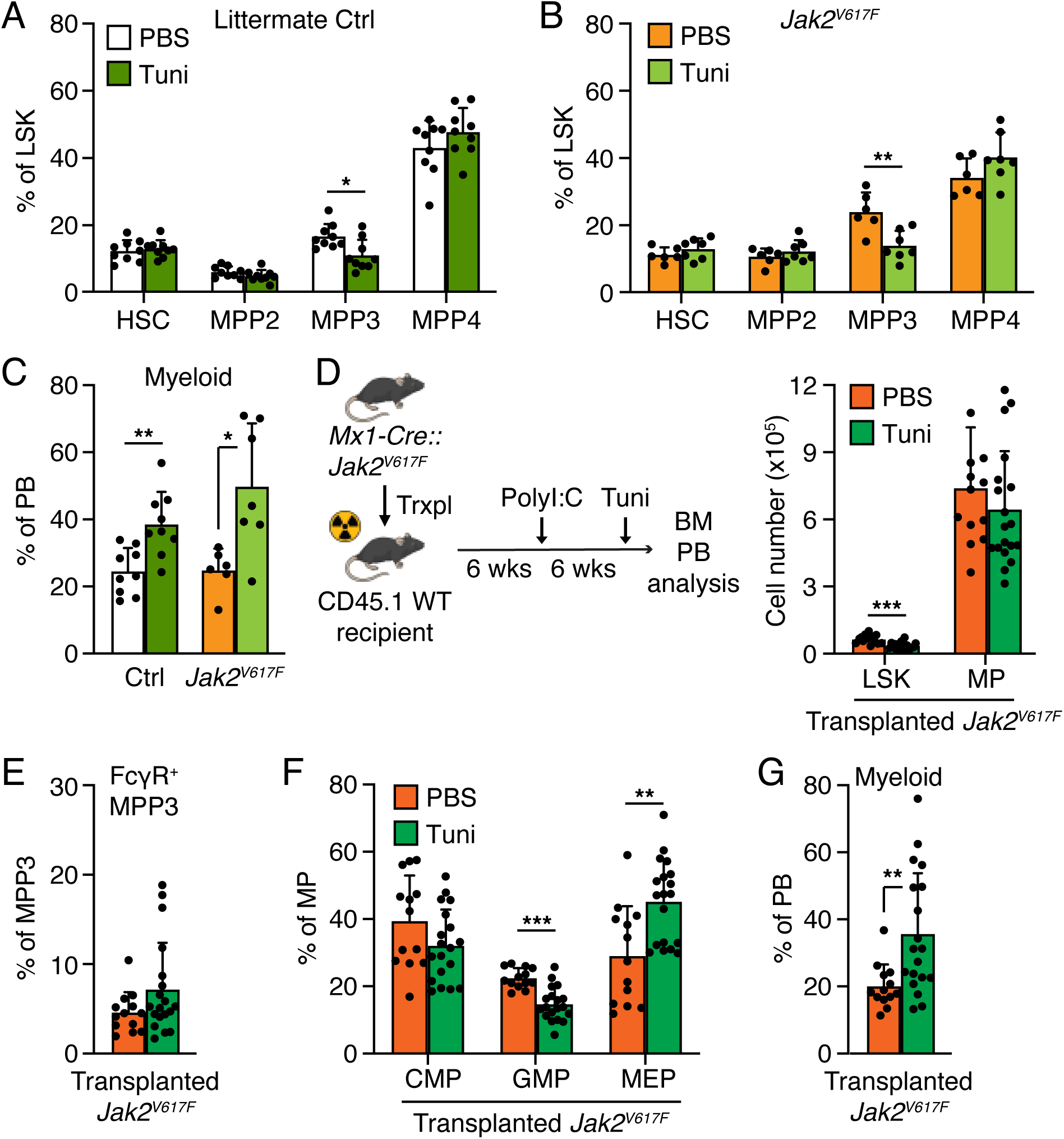
UPR signaling acts at multiple levels of the hematopoietic hierarchy to promote myeloid cell production. (A and B) Hematopoietic stem and progenitor cell (HSPC) population changes in littermate control (Ctrl) (A) and *Jak2^V617F^* mice (B) upon tunicamycin (Tuni) treatment. Results are expressed as a percentage of LSK. (C) Quantification of peripheral blood (PB) myeloid cells upon tunicamycin treatment in littermate controls and *Jak2^V617F^* mice. (D) Experimental scheme (left) and LSK and MP population changes in transplanted *Jak2^V617F^* mice upon tunicamycin treatment (right). One million bone marrow (BM) cells from CD45.2 *Mx1-Cre::Jak2^V617F^*(*Jak2^V617F^*) mice were transplanted into lethally irradiated WT recipient mice. Six weeks post-transplantation, 10 μg polyI:C was injected three times every other day to induce *Jak2^V617F^* expression. After six-week induction, 100 ng/g tunicamycin was injected for five consecutive days, and BM populations and PB myeloid cells were analyzed. Trxpl, transplantation; WT, wild type; Tuni, tunicamycin. (E-G) Changes in FcγR^+^ MPP3 (E), myeloid progenitor (MP) population (F), and PB myeloid cells (G) in transplanted *Jak2^V617F^*mice upon tunicamycin injection. Data are means + S.D., and statistical significance was assessed by a two-tailed unpaired Student’s *t-test*. * *p* ≤ 0.05, ** *p* ≤ 0.01, *** *p* ≤ 0.001.

## Notes

### Summary of Updates

We added more mechanistic data, more biological replicates, more details in the methods and figure legends, and a new supplemental table. We also added a new author.

